# Social experience influences thermal sensitivity: lessons from an amphibious mangrove fish

**DOI:** 10.1101/2023.02.12.528202

**Authors:** Chloé A. Melanson, Claire Allore, Simon G. Lamarre, Suzanne Currie

## Abstract

Understanding factors affecting ectothermic fishes’ capacity to cope with warming temperature is critical given predicted climate change scenarios. We know that a fish’s social environment introduces plasticity in how it responds to high temperature. However, the magnitude of this plasticity and the mechanisms underlying socially-modulated thermal responses are unknown. Using the amphibious, selfing hermaphroditic mangrove rivulus fish (*Kryptolebias marmoratus*) as a model, we tested three hypotheses: 1) social stimulation affects physiological and behavioural thermal responses of isogenic lineages of fish, 2) social experience and acute social stimulation result in distinct physiological and behavioural responses, and 3) a desensitization of thermal receptors is responsible for socially modulated thermal responses. To test the first two hypotheses, we measured the temperature at which fish emerged (i.e., *pejus* temperature) with acute warming with socially naïve, isolated fish and socially experienced fish. Our results did not support our first hypothesis as fish socially-stimulated by mirrors during warming (i.e., acute social stimulation) emerged at similar temperatures as isolated fish. However, in support of our second hypothesis, prior social experience resulted in fish emerging at a higher temperature than socially naïve fish suggesting an increase in *pejus* temperature with social experience. We measured whole-body cortisol concentrations of socially naïve and socially experienced fish and determined that socially experienced fish had significantly higher cortisol concentrations than socially naïve fish. To test our third hypothesis, we exposed socially experienced and naïve fish to capsaicin, an agonist of TRPV1 thermal receptors. Socially experienced fish emerged at significantly higher capsaicin concentrations than socially naïve fish suggesting a desensitization of their TRPV1 thermal receptors. Collectively, our data indicate that past and present social experiences impact the behavioural response of fish to high temperature. We also provide novel data suggesting that social experience affects the capacity of fish to perceive warm temperature.

## Introduction

Warming of the oceans can cause species migrations (García-Molinos *et al*., 2016), decline in marine biomass (Lotze *et al*., 2019), higher extinction rates (Wabnitz *et al*., 2018) as well as changes in community structure (MacNeil *et al*., 2010) and ecosystem functions (Petchey *et al*., 1999). Therefore, predicted ocean warming will have critical biotic impacts (Madeira *et al*., 2018), especially on ectotherms, who rely largely on their environment to thermoregulate. The physiological response of ectotherms to warming can be illustrated with the help of thermal performance curves (TPC) (Schulte *et al*., 2011; Sinclair *et al*., 2016). TPCs model how an ectotherm’s body temperature affects its instantaneous performance and define the temperature interval in which a species can survive (Krenek *et al*., 2011). In these theoretical curves, the animal’s *pejus* temperature lies outside the optimal thermal range and is the temperature at which physiological processes cease to be optimal and “get worse” (Pörtner, 2010). Organisms can survive temperatures beyond their *pejus* temperature thanks to physiological mechanisms such as the heat shock response, metabolic depression, and antioxidative defence, but they can only do so for a limited time (Pörtner *et al*., 2017). Thus, it is crucial that we understand the factors that can affect the *pejus* temperature of tropical marine species to understand their thermal limits.

The impact of temperature depends not only on the degree of warming but also on the susceptibility of species to warming (Deutsch *et al*., 2008; Tewksbury *et al*., 2008). This susceptibility, defined by the width of the TPC, can be influenced by a variety of factors including genetics (Manis & Claussen, 1986), age (Bowler & Terblanche, 2008), gender (Sornom *et al*., 2010), body size (Cheung *et al*., 2012; Peck *et al*., 2009) recent thermal history (Galbraith *et al*., 2012) and, important for this study, sociality (LeBlanc et al., 2011). For example, social signals disrupted the adaptive thermal response of wild, tropical mangrove rivulus (*Kryptolebias marmoratus*) causing these amphibious fish to emerge from water, a marker of their *pejus* temperature, at higher temperatures than their isolated counterparts (Currie and Tattersall 2018). In gregarious juvenile lake sturgeon (*Acipenser fulvescens*), the presence of conspecifics during acute thermal exposure caused a decrease in both endocrine and cellular stress responses (Yusishen *et al*., 2020). The presence of conspecifics has also been shown to modify the preferred and threshold temperatures of the black-axil chromis (*Chromis atripectoralis*) (Nay *et al*., 2021). Not only can acute social stimulation affect a fish’s response to temperature, but social experience from prior social stimulation may also be influential. Indeed, social experience can affect an individual’s basal cortisol concentrations and neurophysiological systems and, in turn, affect behavioural traits (*Neolamprologus pulcher*, Antunes *et al*., 2021a; Antunes *et al*., 2021b). Despite the growing evidence that social stimulation affects responses to temperature in fishes, the nature of the social interactions and the mechanism underpinning these responses remain unclear.

To react to changing temperature, animals first need to be able to perceive or sense temperature. We know that thermosensation depends on the activation of gated ion channels in the nervous system and the skin (Caterina *et al*., 1997). Most of these thermal receptors belong to the transient receptor potential (TRP) ion channel family (Castillo *et al*., 2018) and are highly conserved across taxa (Saito & Shingai, 2006; Saito *et al*., 2011). Six thermosensitive TRP channels have been identified in teleosts and two (TRPV1 and TRPV4) help to perceive ambient environmental temperature (Saito & Shingai, 2006; Saito *et al*., 2011). All of these thermoTRP channels have distinct temperature intervals at which they are activated (Islas, 2017). Although TRPV1 is the best-studied thermoTRP channel, we have limited knowledge of its mechanism of activation in any organism. In addition, its function varies across species which means we must generalize cautiously (Caterina *et al*., 1997, Saito & Shingai, 2006; Garcí-Ávila & Islas, 2019). It is believed that TRPV1 is activated at temperatures exceeding 42 °C (Caterina *et al*., 2000; Caterina, 2007), but its activation temperature in fish seems to be lower, at ~ 32-33 °C (zebrafish *Danio rerio*, Gau *et al*., 2013; rainbow trout *Oncorhynchus mykiss*, Ashley *et al*., 2007), although there are limited data available. Here, we hypothesized that an individual’s ability to sense temperature could be influenced by its social environment. If socially-modulated thermal risk, as observed in wild mangrove rivulus who delay their emersion out of critically warm water (Currie & Tattersall, 2018), could be explained by variation in thermosensation, then changes in the social environment would result in differences in thermal sensitivity.

We used isogenic lineages of the tropical mangrove rivulus in different social situations to determine the possible effects of sociality on thermal biology. In general, the *pejus* temperatures of tropical marine ectotherms’ are closer to their critical thermal maximum (CT_max_) than their temperate counterparts, making them particularly vulnerable to warming temperatures (Sunday *et al*., 2011; Huey *et al*., 2009; Krenek *et al*., 2012). The mangrove rivulus is a simultaneous hermaphrodite fish and is capable of producing fertilized eggs (Taylor *et al*., 2001). Since genetic profiles and relatedness can influence how individuals interact (Nakamura *et al*., 2016), studying multiple isogenic lineages allowed us to investigate the effect of genetics on sociality (Hsu *et al*., 2008). We used socially naïve and socially experienced fish that were isolated or acutely socially stimulated by a mirror or conspecifics. We measured key behaviours to assess whether or not the different responses to warming were due to varying levels of aggression or interest towards the social stimulus. Our overarching hypothesis was that the social environment modulates physiological and behavioural thermal responses within and between isogenic lineages of fish due to different thermosensing capacities. We capitalized on the amphibious nature of mangrove rivulus and the fact that it is a self-fertilizing hermaphrodite and first predicted that socially stimulated fish (fish with a mirror or conspecifics) would emerge from water at a higher temperature than isolated and socially naïve fish. In an attempt to understand the mechanism underpinning socially-influenced thermal responses, we hypothesized that the social environment would result in a desensitization of thermoreceptors. To this end, we predicted that socially experienced fish treated with the well-known TRPV1 agonist, capsaicin, would require higher concentrations to elicit a thermal escape response (i.e., emersion) compared to socially naïve fish, suggesting a desensitization of these thermal receptors.

## Materials and Methods

### Experimental Animals

We performed all experiments with adult mangrove rivulus hermaphrodites (*K. marmoratus*) housed in a breeding colony at the Acadia University Animal Care Facility, Wolfville, NS, Canada. The three isogenic lineages in this colony, Belize1 (50.91), Belize2 (Dan06), and Honduras (Hon11), were originally caught in Twin Cayes, Belize in 1991, in Dangriga, Belize in 2006 and in the Bay Islands, Honduras, in 1996, respectively (Tatarenkov *et al*., 2010). The three lineages are genetically divergent; each strain has been bred in a laboratory for more than 45 generations and has multiple loci differences (Tatarenkov *et al*., 2010). All fish were adults older than 3 months and reared individually in 120 ml cups of synthetic seawater made with Instant Ocean (Pets and Pond Canada) and reverse osmosis water. The cups were held in constant conditions (80 mL of 15ppt water, 12h: 12h light: dark cycle, 30-75% humidity). We performed water changes biweekly and maintained the colony on a natural thermal diel cycle of 25°C to 28.5°C (Ellison *et al*., 2012). We fed fish three times per week with live *Artemia* sp. nauplii; frozen bloodworms supplemented the fish’s diet once per week. The Acadia University Animal Care Committee approved all protocols in this study (#05-19 & #02-22).

### Experimental Protocol

We provided both mirrors and live conspecifics to study whether or not the source of social stimulation had an effect. We use “social stimulation” to describe an event in which fish respond to a perceived (mirror) or live conspecific and “social experience” to describe a fish that was previously socially stimulated for 24 h.

#### Social experience

Socially naïve fish were raised in isolation and had never interacted with conspecifics prior to the experiments. For socially experienced fish, we placed focal fish in a cylindrical container with a diameter of 11.2 cm filled with 300 ml of 15 ppt salt water. The container was divided into three equal chambers with fiberglass mesh that allowed the fish to interact visually and chemically with two other fish while preventing physical interaction. Because the container was cylindrical, all three fish could equally interact with each other regardless of which chamber we placed them in. Therefore, we randomly assigned chambers to the fish. We habituated the fish to the presence of their size-matched conspecifics for 24 ± 1 h before transferring them to the experimental container for the start of the experiment. Socially naïve fish were not manipulated prior to their transfer to the experimental container. Preliminary trials indicated that manipulation prior to transfer to the experimental container did not result in physiological stress, as determined by whole-body cortisol levels (Fig. S1).

#### Series 1 – Social stimulation impacts physiological and behavioural thermal responses

Here, we tested the hypothesis that social stimulation alters behavioural and physiological thermal responses in three distinct laboratory lineages of mangrove rivulus. We measured the emersion temperature (T_em_: temperature at which the gills of the fish emerged out of water), the critical thermal maximum (CT_max_: temperature at which the fish could not maintain dorsal-ventral equilibrium; Cox, 1974; Beitinger *et al*., 2000; Gibson *et al*., 2015) and the thermal safety margin (TSM: difference between CT_max_ and Tem; Currie and Tattersall 2018) to test this hypothesis. We interpret T_em_ as the *pejus* temperature or T_*pej*_, the water temperature beyond the optimal temperature (T_opt_) when performance “gets worse” and the fish escapes (Pörtner, 2010).

We measured T_em_ and CT_max_ of socially naïve fish in isolation (control) and with a mirror from all three lineages (Fig. 1; Belize1: naïve control n = 16, naive mirror n = 20; Belize2: naïve control n = 13, naïve mirror n = 12; Honduras: naïve control n = 20, naïve mirror n = 20). We analyzed the behaviour of the fish *post-hoc* with video recordings (see *Analysis*).

**Fig. 1.**
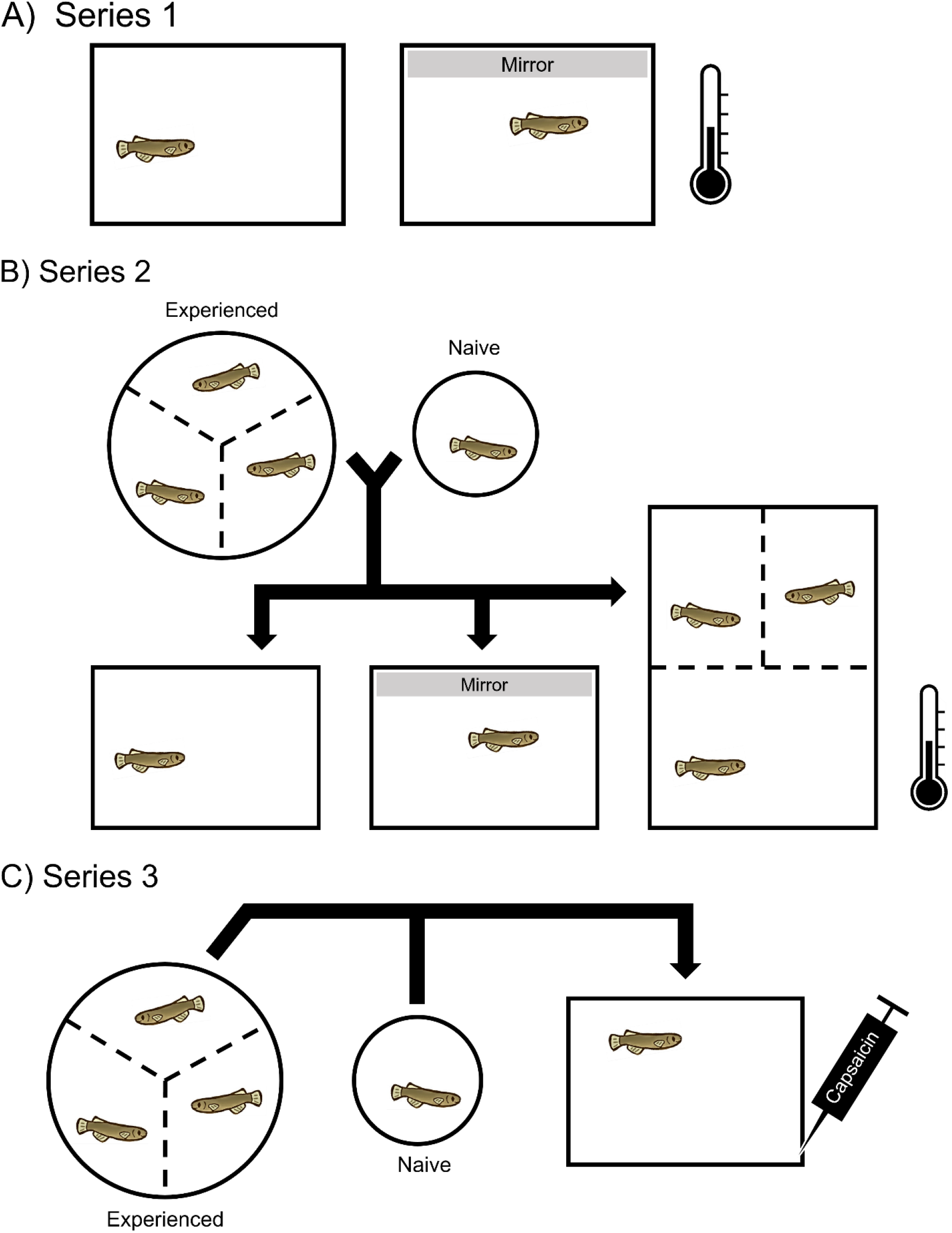
Schematic protocols or Series 1, Series 2 and Series 3. A) was performed with all three lineages, Belize1, Belize2 and Honduras, while B) and C) were only performed with the Honduras lineage.

#### Series 2 – Social experience impacts physiological and behavioural thermal responses

Unlike wild mangrove rivulus (Currie and Tattersall 2018), laboratory lineages did not respond to their mirror reflection in terms of thermal behaviour (see Series 1 Results). Because wild fish are not socially naïve, we hypothesized that social experience affects the response to temperature. To address this second hypothesis, we used one lineage (Honduras) and compared socially naïve and socially experienced fish that were isolated during the acute thermal stress (naive n = 20; experienced n =19; Series 2a). We then compared socially naïve and socially experienced fish that were stimulated by a mirror during the acute thermal stress (naive n = 20; experienced n = 22; Series 2b). In a third experiment, we replaced the mirror with two live size-matched conspecifics (Series 2c). We performed this experiment in a different chamber than Series 2a and Series 2b due to the addition of the two conspecifics (Fig. 1). We measured T_em_ in all three experiments and measured CT_max_ in Series 2a and 2b. Since CT_max_ did not differ between socially naïve and socially experienced fish in Series 2a and 2b, we did not measure it in Series 2c. To determine if the social conditions resulted in physiological stress, we measured whole-body cortisol levels of socially naïve and socially experienced fish before exposure to a thermal stress. We analyzed the behaviour of the fish *post-hoc* with recordings of the experiments (see *Analysis*).

#### Series 3 – Socially modulated response to temperature is the result of desensitization of TRPV1 channels

Here, we measured emersion of socially naïve and experienced fish but instead of increasing the temperature, we used increasing concentrations of the specific TRPV1 agonist, capsaicin (Caterina *et al*., 1997) to elicit chemical warming. We used fish from the Honduras lineage for these experiments. We measured the emersion concentration (C_em_: the concentration of capsaicin at which the gills of the fish emerged out of water), the number of times the fish attempted to emerge from the water as well as the critical concentration maximum (CC_max_: the concentration of capsaicin at which the fish loses equilibrium), if and when it occurred. We made these measurements on both socially naïve and experienced fish (Fig. 1) and had control groups that were treated with water or 0.1% DMSO, capsaicin’s vehicle (naïve water n=10, naïve DMSO n=10, naïve capsaicin n=13, experienced water n=10, experienced DMSO n=10, experienced capsaicin n=13).

### Experimental Analyses

We performed all experiments between 12-5 pm and we recorded them using a camcorder (Canon VIXIA HF200) for *post hoc* behavioural analyses. Whenever we removed a fish from the experimental container, we returned it to its individual cup which held 80 ml of 15 ppt water at 32-34 °C and then to the colony. We measured the length (to nearest mm) and mass (to the nearest mg) of fish at least 24 h after they were subjected to a test. When we used focal fish in more than one experiment, the time interval between experiments was at least 4 months.

#### Emersion temperature (T_em_, Series 1 & 2)

T_em_ experiments started with a 1 h familiarization in the new chamber. During this time, we kept the chamber in a 25 −28 °C water bath and then heated the chamber at a rate of 1 °C · min^-1^ in a water bath (VWR, model: WBE05) containing 2 L of water. The experiment ended when the gills of the fish emerged from the water (T_em_) (Gibson *et al*., 2015; Currie & Tattersall, 2018). In all T_em_ measurements, we used a steel temperature probe connected to a Lab Pro (Vernier Software & Technology) to measure the water temperature to the tenth of a degree every second.

For T_em_ measurements in which the fish was isolated or had access to a mirror, we placed the focal fish in one of four identical and visually isolated experimental chambers (8L x 4W x 7H cm^3^) filled with 150 ml of 15 ppt water. If the experiment required social stimulation, we removed an opaque barrier against the long side of the chamber to reveal either a mirror (Series 1 & Series 2b) or two live conspecifics (Series 2c). We revealed the mirror at the beginning of heating and revealed the two stimulus fish 10 min before the start of the heating; this ensured that the focal fish perceived them and was socially stimulated throughout the acute warming. A fiberglass mesh barrier allowed focal and stimulus fish to interact visually and chemically but not physically. The focal fish were unfamiliar to the two stimulus fish and all fish were of the same lineage. To closely observe the focal fish, we carefully removed both stimulus fish from the experimental container when the water reached 40 ° C (close to CT_max_) and returned them to their individual cups. When we allowed the fish to interact with two live conspecifics, the focal fish chamber (8 L x 4W x 7H cm^3^) held 150 ml of 15 ppt salt water while the two stimulus fish chambers (each 4L x 4W x 7H cm^3^) on the other side of the fiberglass mesh held 75 ml each of 15 ppt salt water.

#### Critical thermal maximum (CT_max_, Series 1 & 2)

We measured CT_max_ in a way similar to T_em_, but with a few modifications. We filled the chamber used in Series 1, Series 2a, and Series 2b with 190 ml of water and covered with cellophane to prevent the fish from emerging as the water warmed. We ended the experiment when the fish could not maintain dorsal-ventral equilibrium (Currie & Tattersall, 2018).

We measured CT_max_ on the same fish as we measured T_em_, but performed the second measurement 48 h after the first. To evaluate whether the order of testing had an impact on the results, we alternated the order in which both tests were performed.

#### Thermal safety margin calculations (TSM, Series 1 & 2)

We calculated TSM by using the following formula (Currie & Tattersall 2018):

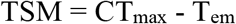

#### Does social stimulation result in physiological stress? (Series 2)

In Series 2, we investigated the effect of social experience on the behavioural response to warm temperatures. Following the 24 h experience period when we exposed fish to conspecifics or kept them isolated, the fish was then sacrificed in an ice water bath, homogenized and we measured whole-body cortisol concentration. The homogenization buffer was in mM: 81 Na_2_HPO_4_, 24.7 NaH_2_PO_4_, 99.9 NaCl, 1 EDTA and we used a tissue grinder (Fisherbrand Pellet Pestle). We extracted cortisol following Ridgeway *et al*., (2021) except we used an enzyme-linked immuosorbent assay (ELISA) (Product #402710, Neogen). The plates were read at 450 nm in a SpectraMax M5 with SpectraMax software.

#### Emersion concentration (C_em_, Series 3)

At the beginning of each experiment, we placed the fish in one of two identical and visually isolated experimental chambers (8L x 4W x 7H cm^3^) filled with 150 ml of 15 ppt water and surrounded by a 2 cm wide ledge and a 10 cm high wall. We placed a stir bar at the bottom of each chamber, which was separated from the fish by a 1.3 cm tall fiberglass mesh insert. We set the stirring speed so that the water in the chamber was mixed homogeneously in approximately 20 s. We kept the chamber at 26.5 ± 2.5 °C with one of two stirring hot plates (Thermolyne, model: SP18425; Fisher Scientific, model: SPN105082). After a 1 h familiarisation period with the chamber, we chemically heated the water at a rate of 20 μM · min^-1^ by gently pipetting 480 μl of 3.2 mM capsaicin at one of the two corners of the experimental chamber, depending on the location of the fish. The capsaicin solution was added every 30 s to a maximum of 200 μM capsaicin (similar to Endo *et al*., 2020 in zebrafish). Since few studies have exposed fish to waterborne capsaicin, we performed preliminary experiments to determine the capsaicin concentration that would cause emersion (200 μM), but not loss of equilibrium or mortality (see Supplemental information). The experiment ended when the fish reached CC_max_ or 15 min after the start of the experiment, whichever came first. We prepared a stock solution of 195 mM capsaicin in DMSO and the working solutions of 3.2 mM capsaicin was prepared daily by diluting the stock solution in 15 ppt water. We repeated the same protocol with 0.1% DMSO in 15 ppt water instead of a capsaicin solution in the control experiments.

#### Video Analysis of Behaviour

We tracked behaviour through *post hoc* video analysis of 89 experiments in Series 1 and 2 (control naive n=13; control experienced n=15; mirror naive n=17; mirror experienced n=15; fish naive n=14; fish experienced n=14) using EthoVision XT ver. 15.0 (Noldus Information Technology). The experiments were analyzed from start to finish (approx. 15 min). We recorded six behaviours: i) the latency to resume normal activity, ii) latency to first approach the interaction zone (defined as one body length of the individual, starting at the barrier between the social stimulus and the focal fish), iii) total time spent in the interaction zone (association time), iv) time spent swimming, v) time in contact with the social stimulus barrier (Li *et al*., 2018; Edenbrow & Croft, 2013), and vi) the frequency of aggressive approaches towards the social stimulus (fish or mirror). For this latter behaviour, we manually measured it (control naive n=13; control experienced n=15; mirror naive n=16; mirror experienced n=15; fish naive n=14; fish experienced n=13). To measure behaviour elicited by the social stimulus alone, without warming, we visually analyzed 5 min at the start of each experiment starting at 1:00 min to ensure that the fish had resumed normal activity. We defined aggressive approaches as attempted bites and mouth wrestling against the social stimulus barrier, swimming in bursts along or towards the social stimulus barrier and lateral displays performed parallel to the social stimulus barrier. Frequency of aggressive approaches is included in association time given that we measured time spent in the interaction zone (i.e., association time) independently of behaviour. We analyzed these six behavioural markers of aggression and activity to possibly explain differences between socially stimulated fish and isolated fish as well as between socially experienced and socially naïve fish. We also compared behavioural markers of fish with different sources of social stimulation (i.e., mirror vs live conspecifics).

### Statistical Analysis

We performed all statistical analyses using RStudio (v. 4.1.0) and used GraphPad Prism 9.2.0 (332) for Series 3, all with a critical alpha value of 0.05. We used Shapiro-Wilk and Bartlett’s tests to test the assumptions of normality and homoscedasticity, respectively.

In Series 1, we performed three-way analyses of covariance (ANCOVA) with the independent variables of lineage and treatment (control or mirror) and the dependent variables: T_em_, CT_max_ and TSM. Fish condition index, mass and length were covariates. In Series 2, we performed two-way ANOVAs to compare the T_em_ of naïve and experienced fish of Series 2a and 2b. The independent variables of the two-way ANOVAs were social experience and treatment (control or mirror) and the the dependent variables: T_em_, CT_max_ and TSM. The residuals of Series 2’s two-way ANOVA respected the assumption of normality, but deviated slightly from the assumption of homoscedasticity. We judged the test to be acceptable despite the modest deviation from homoscedasticity. We used a t-test to compare the T_em_ of naïve and experienced fish with live conspecifics (Series 2c). We used a t-test to determine if there was a significance in cortisol concentration between socially naïve and socially experienced fish. We also used t-tests to compare association time and non-parametric Mann-Whitney tests to compare aggressive approach frequency between socially naive and socially experienced fish. We computed non-parametric Spearman coefficients to study the relationship between association time and T_em_ and to analyze the relationship between aggressive approach frequency and T_em_ of socially naïve and socially experienced fish.

To compare the capsaicin response of socially naïve and socially experienced fish in Series 3, we used dose-response curves and compaired the half maximal effective concentration (EC50) of capsaicin of socially naive and socially experienced fish using a modified extra-sum-of-squares F test.

## Results

### Series 1 – Social stimulation impacts physiological and behavioural thermal responses

In socially naïve fish from three isogenic lineages, the social stimulus provided by the mirror did not significantly affect T_em_ when compared to fish with no mirror (Table S1; Fig. S2). The social stimulus provided by the mirror also had no significant effect on CT_max_ (Table 1; Table S2).

**Table 1.** Thermal responses *K. marmoratus* of Belize1, Belize2 and Honduras lineages.

|  | **T<sub>em</sub> (°C)** |  | **CT<sub>max</sub> (°C)** |  | **TSM (°C)** |  | **N** |  |
| --- | --- | --- | --- | --- | --- | --- | --- | --- |
|  | **Control** | **Mirror** | **Control** | **Mirror** | **Control** | **Mirror** | **Control** | **Mirror** |
| **Belize1** |  |  |  |  |  |  |  |  |
| **Naive** | 13.02<br>± 0.14 | 13.02<br>± 0.12 | 39.97<br>± 0.13 | 40.09<br>± 0.13 | -0.05 ±<br>0.19 | 0.07 ±<br>0.17 | 16 | 20 |
| **Belize2** |  |  |  |  |  |  |  |  |
| **Naive** | 12.94<br>± 0.19 | 12.76<br>± 0.20 | 40.08<br>± 0.14 | 40.10<br>± 0.15 | 0.15<br>± 0.22 | 0.34<br>± 0.19 | 13 | 12 |
| **Honduras** |  |  |  |  |  |  |  |  |
| **Naive** | 13.10<br>± 0.21 | 13.37<br>± 0.13 | 40.20<br>± 0.13 | 40.05<br>± 0.13 | 0.10<br>± 0.21 | -0.32<br>± 0.15 | 20 | 20 |
| **Experienced** | 13.71<br>± 0.11* | 13.81<br>± 0.09* | 40.27<br>± 0.20 | 40.16<br>± 0.15 | -0.43<br>± 0.21 | -0.65<br>± 0.20 | 19 | 22 |

Values are presented as mean ± SEM.
Fish were either not provided a social stimulus during the experiment (control) or had access to a mirror during the experiment (mirror). The fish were subjected to a heating protocol of 1.0 °C·min<sup>-1</sup>.
\*Significant difference (p < 0.05) from naïve

Given that each lineage is isogenic, we were interested in the intraspecific variation in each of our conditions as a gauge of trait plasticity. When comparing across lineages, we found no significant differences in T_em_ (Table S1), CT_max_ (Table S2) or TSM (Table S3). Mass and length of fish did not influence T_em_, CT_max_ or TSM. However, Fulton’s condition factor (k) did affect CT_max_; its effect varied between social stimuli (control or mirror) and among lineages (Table S2). In Belize1 and Belize2, CT_max_ was positively correlated to the condition factor while it was not the case in the Honduras lineage. In addition, the variability of T_em_, CT_max_ or TSM did not differ among lineages. Within lineages, only Honduras fish showed higher variability when isolated compared to fish faced with a mirror (Table S4).

### Series 2 – Social experience impacts physiological and behavioural thermal responses

As with the socially naïve mangrove rivulus, the T_em_, CT_max_ and TSM of socially experienced mangrove rivulus did not differ whether they had access to a mirror or not (Table 1). However, T_em_ was significantly higher in socially experienced mangrove rivulus compared with socially naïve fish. This was only true during control experiments when fish were not socially stimulated (Fig. 2A, Table S4). Even though mirror reflections masked the effect of social experience, we did not observe a significant interaction between social experience and social stimulation. Furthermore, when we provided a social stimulus in the form of live conspecifics during the thermal stress, the effect of social experience on the T_em_ observed with control fish also disappeared (Fig. 2B; t_(df=29)_=1.495, p=0.1457).

**Figure 2.**
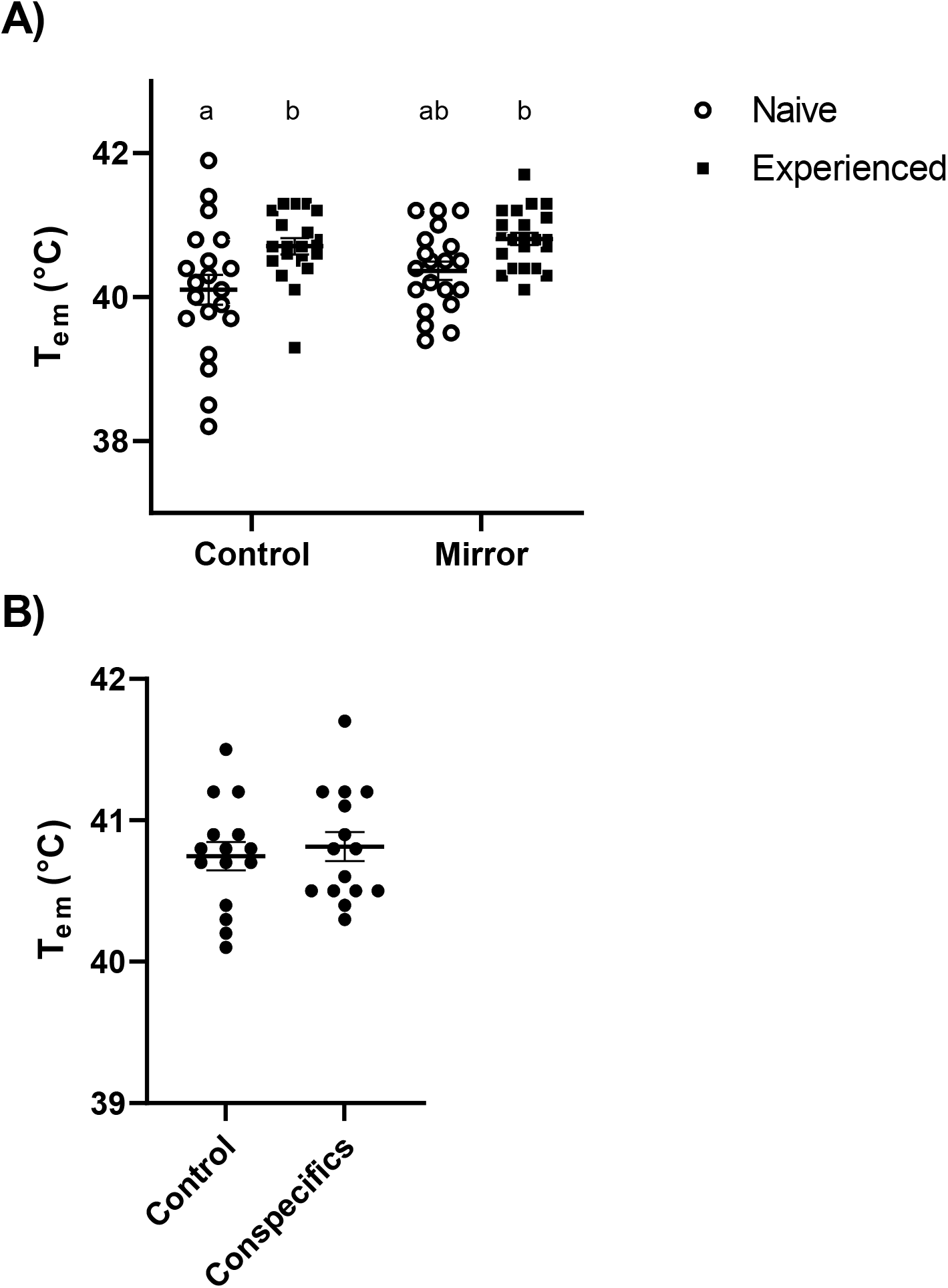
Mean T_em_ (± SEM) of *K. marmoratus* from the Honduras strain. (A) Socially naïve and socially experienced fish had either no social stimulation (Control) or access to a mirror during the experiment (Mirror). (B) Naïve fish were also given access to two size-matched conspecifics during the experiment (Conspecifics). Fish were subjected to a heating protocol of 1.0 °C.min^-1^. There was a significant difference between all socially naive and all socially experienced fish (A, see Table S5). A) Control Naive N= 20; Control Experienced N= 19; Mirror Naive N=20; Mirror Experienced N=22; Table S5. B) Naive N=15; Experienced N=16; p=0.1457.

Despite differences in T_em_, there was no significant difference between the CT_max_ or TSM of socially naïve fish and socially experienced fish (Table 1). Variability of T_em_, CT_max_ and TSM did not differ between socially naïve and experienced fish except for T_em_ of fish with mirror reflections (Table S6). It is noteworthy that some of the T_em_ measurements were beyond the CT_max_ of the fish, whether they were socially naïve, socially experienced, isolated or faced with a mirror. This was the case for all three lineages (Fig. S4).

We compared six behavioural markers of aggression and association between control, mirror and fish treatment groups of the Honduras strain. Since aggression and sociality are not always correlated (Lacasse & Aubin-Horth, 2014), it was important not to exclusively record markers of aggression. The association time for focal fish did not differ between socially naïve and socially experienced fish when they were isolated during the experiment (Fig. 3A) or when they had access to a mirror (Fig. 3B). However, when the fish were with live conspecifics during the experiment, socially naïve fish spent significantly more time interacting with them than socially experienced fish (Fig. 3C). The association time was not significantly correlated with T_em_ in any experimental group except for socially experienced fish faced with a mirror (Fig. S5). The frequency of aggressive approaches did not differ between socially naïve and socially experienced fish. This was true when fish were isolated during the experiment, had access to a mirror or could interact with conspecifics (Fig. S3). Aggression was also not correlated with T_em_, whether the fish were socially naïve or experienced or had social stimulation (Fig. S6). Although we did not observe differences in aggression in socially experienced fish with the mirror compared to control (Fig. S3B), the mirror caused the T_em_ difference between socially naive and socially experienced fish to disappear (Fig. 2A, Table S4).

**Figure 3.**
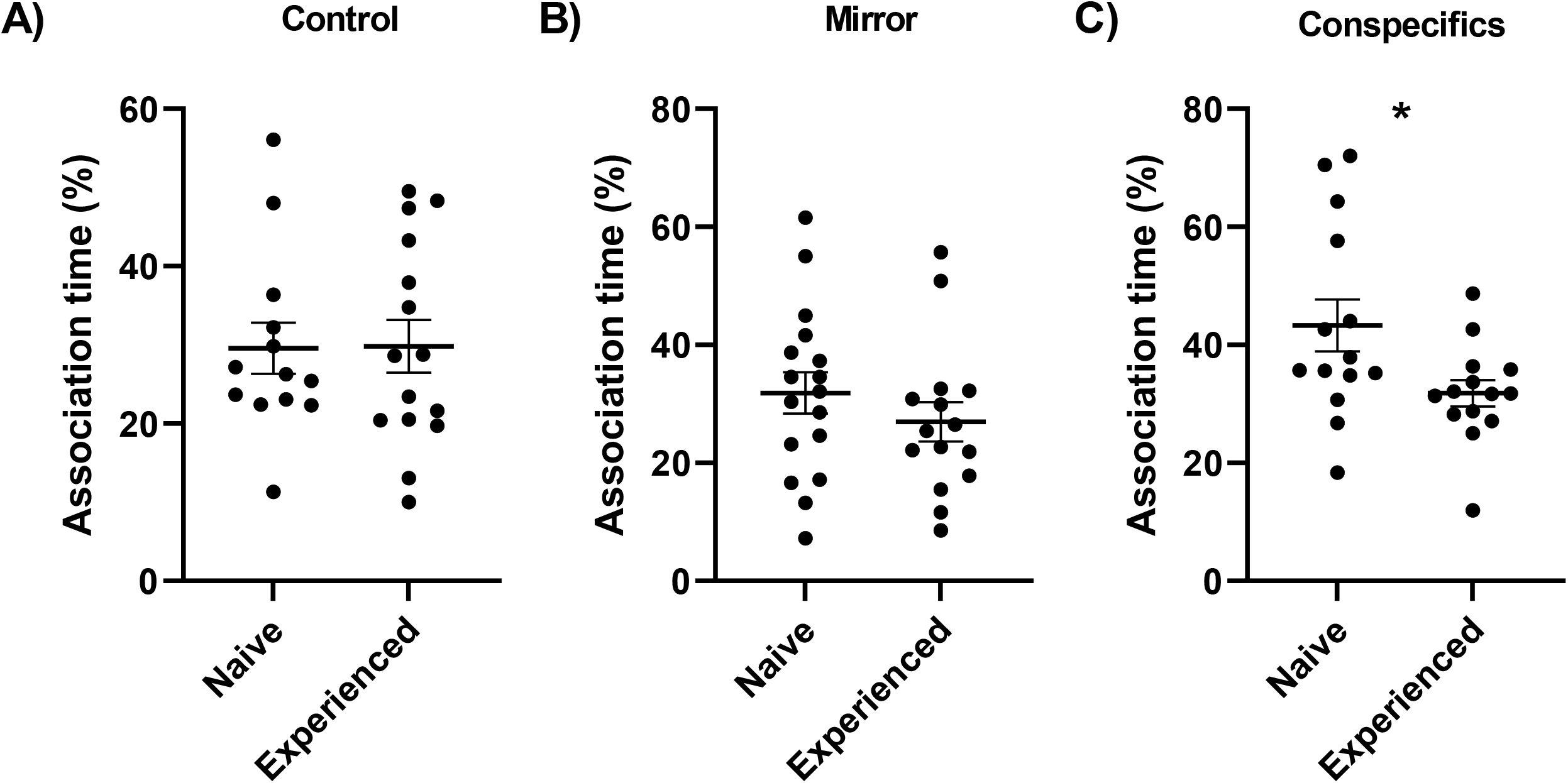
Mean association time with the social stimulus (± SEM) of socially naive and socially experienced *K. marmoratus* from the Honduras strain. (A) Control fish did not have a social stimulus during the experiment, (B) fish had access to a mirror during the experiment, (C) fish were exposed to two size-matched conspecifics during the experiment. Fish were subjected to a heating protocol of 1.0 °C.min^-1^. A) Naive N=13; Experienced N=15; p=0.9551. B) Naive N=17; Experienced N=15; p=0.3222. C) Naive N=14; Experienced N=14; p=0.0285.

To determine if there were differences in physiological stress with social experience, we measured whole-body cortisol concentrations. Socially experienced fish had significantly higher whole body cortisol levels compared to socially naive fish (Fig. 4; t_(df=24)_=2.903; p=0.0078).

**Figure 4.**
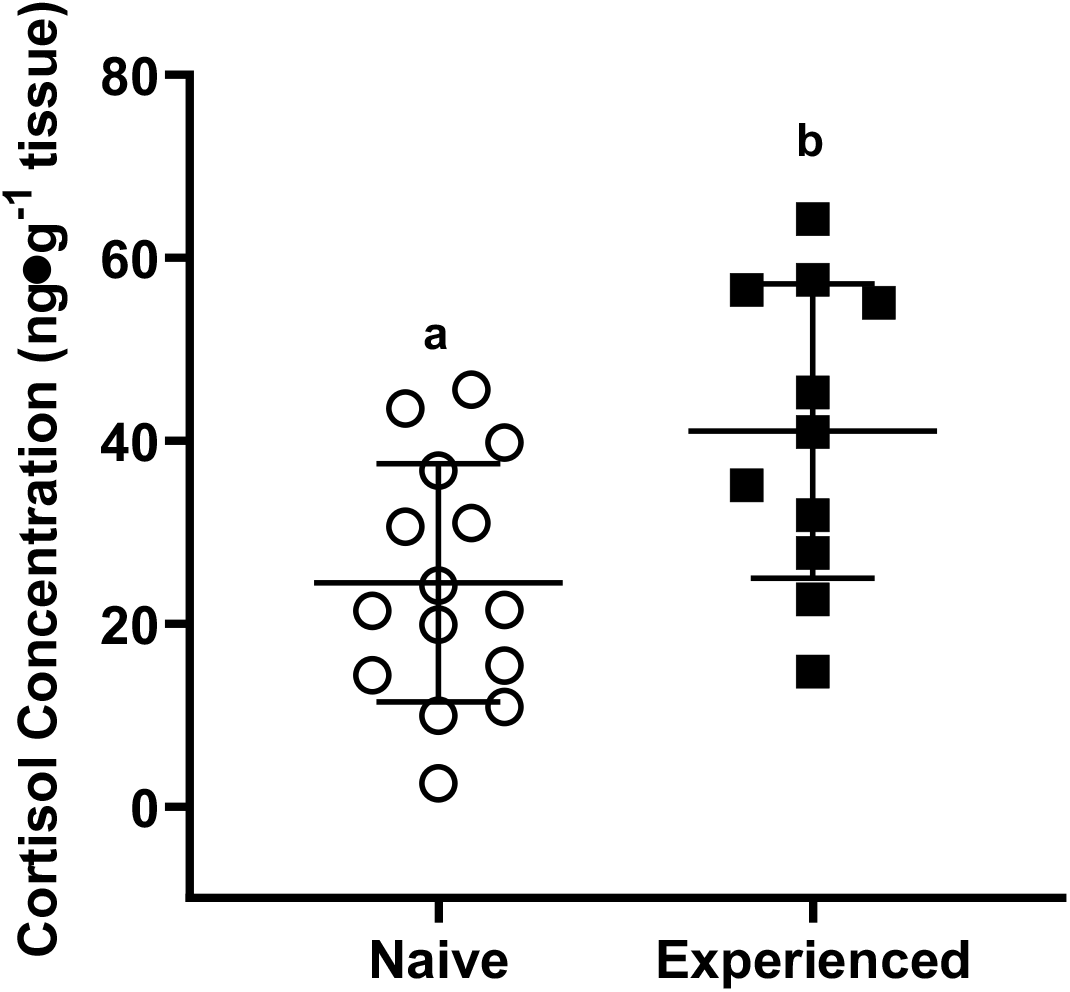
Mean whole-body cortisol levels (ng.g^-1^ tissue) (± SEM) of *K. marmoratus*. Fish were isolated (naive) or subjected to a 24 h interaction period with two size-matched conspecifics prior to the measurement (experienced). Naive N=15; Experienced N=11. Different letters indicate significance (α <0.05), p=0.0078.

### Series 3 – Socially modulated response to temperature is the result of desensitization of TRPV1 channels

We then tested if the desensitization of TRPV1 could provide a mechanism to explain how social experience influences the thermal response of mangrove rivulus. By specifically activating TRPV1 receptors with capsaicin, we were able to test whether or not the difference we observed in T_em_ in different social contexts was because of different thermal sensing capacities. Because we observed a significant correlation between fish mass and capsaicin concentration at first emersion for socially naïve fish (Fig. S7), we normalized the capsaicin concentration per mg of fish (Fig. 5). Socially experienced fish emerged at significantly higher capsaicin concentrations (Fig. 5; naïve EC50=2.921 μM·mg^-1^; experienced EC50=5.286 μM·mg^-1^) compared to naïve fish. This was true whether or not capsaicin concentration was normalized per mg of fish.

**Figure 5.**
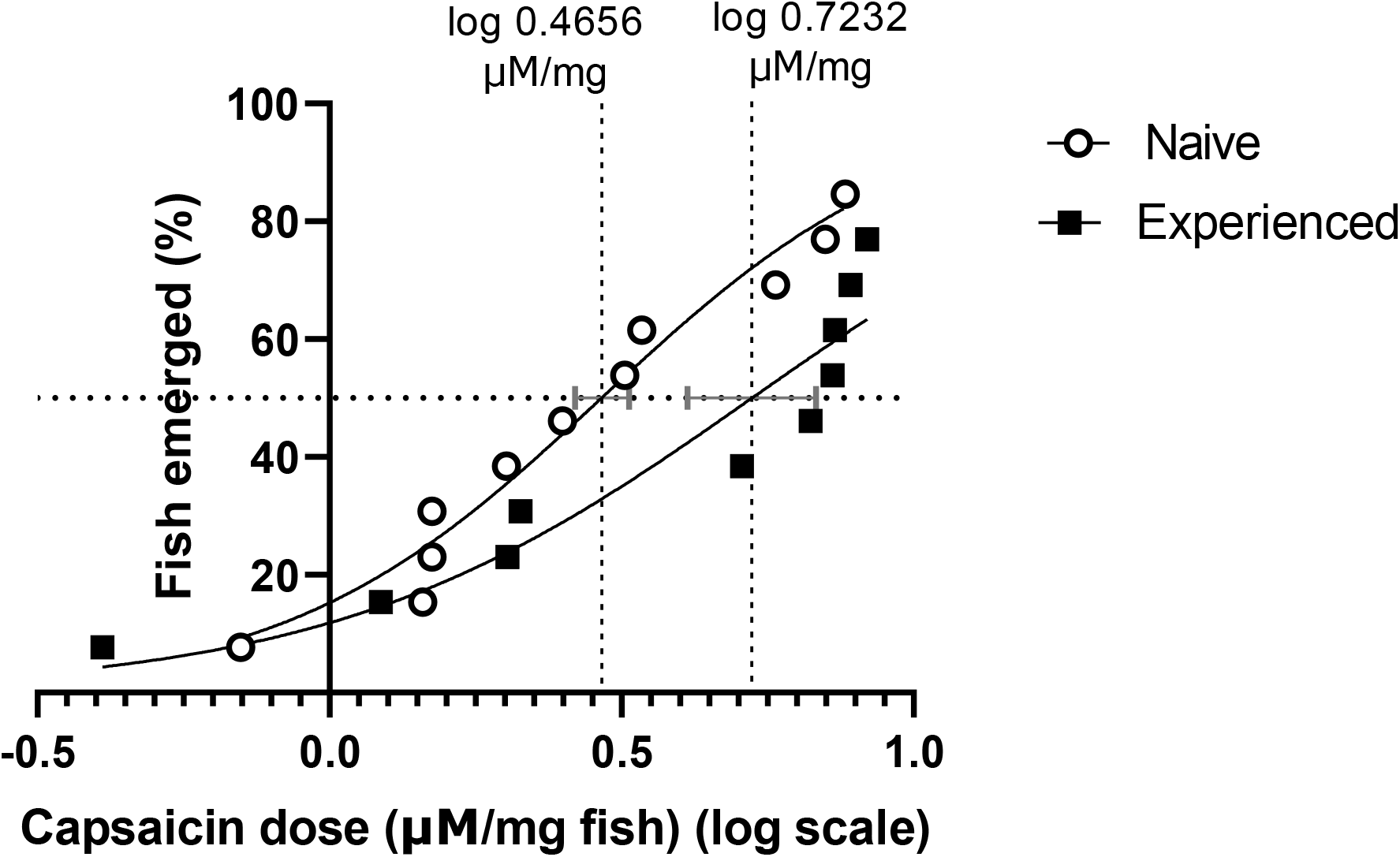
Dose response curve for EC50 analysis. Emersion response of *K. marmoratus* from the Honduras strain to capsaicin concentrations (μM) normalized by mass (mg). Fish were socially isolated until the experiment (naive) or were subjected to a 24 h interaction period with two size-matched conspecifics prior to the experiment (experienced). Naive EC50 = 2.921 μM.mg^-1^; Experienced EC50 = 5.286 μM.mg^-1^. Naive cap N=13; Exp cap N=13.

Whether or not fish were socially naïve or experienced, fish exposed only to water or DMSO emerged significantly less than fish exposed to capsaicin and never reached CC_max_ (Fig. S8A & Fig. S8C). In fact, only one of the forty controls emerged and it did so only once (Fig. S8A). Despite emerging at significantly higher capsaicin concentrations, socially experienced fish and socially naive fish emerged at the same frequency (Fig. S8A).

## Discussion

We tested the hypothesis that social stimulation and experience affect physiological and behavioural thermal responses within and between isogenic lineages of fish due to varying thermosensing capacities. Our results indicate that acute social stimulation with a mirror reflection or with live conspecifics do not affect the thermal response in lab lineages, unlike in wild fish (Currie and Tattersall 2018). However, social experience increased the *pejus* temperature (T_em_) of fish, an effect masked by acute social stimulation (mirror or live fish) thus modulating thermal responses. We present evidence that socially experienced fish have different responses to a TRPV1 agonist compared to naïve fish, suggesting that the mechanism underlying socially influenced thermal responses is linked to thermosensation. Specifically, socially experienced fish emerged at capsaicin concentrations almost twice as high as socially naive fish.

We first stimulated a focal fish by exposing them to a mirror or two live conspecifics to the experimental chamber. We did this with three distinct isogenic lineages in an attempt to determine whether or not there is a genetic component to the response. Because the presence of a mirror delayed the emersion temperature in wild *K. marmoratus* (Currie & Tattersall, 2018), we predicted that our laboratory isogenic fish would exhibit the same delayed emersion with social stimuli from a mirror or with conspecifics. Surprisingly, the presence of a mirror had no effect on the T_em_, CT_max_, or TSM in our three lineages of laboratory *K. marmoratus* nor did the presence of two conspecifics. The absence of significant differences in thermal response among our three isogenic lineages suggests that the socially-influenced thermal responses we were studying are not driven by genetics. Because of this, we performed all subsequent experiments with a single lineage (Honduras). It is worth noting that, unlike our study, Currie and Tattersall (2018) used a paired design where each fish was tested with and without a mirror. However, the most likely reason for the differences between wild and laboratory fish is the source of the fish. Unlike laboratory lineages, wild fish are not socially naïve.

It is well established that wild animals and laboratory animals do not always yield similar results, especially in behavioural and endocrine studies (Calisi & Bentley, 2009). More specifically, laboratory fish have been shown to have different thermal and social responses than their wild counterparts. For example, laboratory-held zebrafish (*Danio rerio*) have reduced thermal plasticity (Morgan *et al*., 2022) and had significantly higher thermal tolerances (CT_max_) than wild zebrafish (Morgan *et al*., 2019) while the opposite was true in trout (*Salmo trutta, Salvelinus fontinalis* and *Oncorhynchus mykiss*, Carline & Machung, 2011). Furthermore, domesticated fish generally have modified aggressive behaviour (Lucas *et al*., 2004; Jonsson & Jonsson, 2006; Campbell *et al*., 2015) and courtship behaviour (Fleming *et al*., 1996). In particular, our laboratory fish are homozygous, whereas wild fish are not (Ellison *et al*., 2012, Tatarenkov *et al*., 2010). Moreover, our *K. marmoratus* had also been raised in isolation and had significantly different social histories than the wild mangrove rivulus, which might also explain why social cues had different effects. Why wild mangrove rivulus responded more strongly to mirrors (Currie & Tattersall, 2018) than our laboratory fish is unknown, but it may be due to genetic differences between our laboratory lineages and the wild populations, different environmental holding conditions and/or different social histories (Mayer *et al*., 2009; Price, 1999).

We know that social experience can affect how fish respond to temperature. For example, the social status of rainbow trout (*O. mykiss*) influenced their heat shock response and thermal tolerance (LeBlanc *et al*., 2011; Bard *et al*., 2021). Convict cichlids (*Amatitlania nigrofasciata*) that had previously socialized with mixed-sex groups were able to create pair bonds with new fish more frequently than fish that only had same-sex social experience (Little *et al*., 2017). In the same species, dyadic contests between individuals housed in isolation lead to more violent interactions (e.g., mouthwrestling and attack-bite sequences) than contests between individuals that had been housed in communal tanks (Earley *et al*., 2006). Given the influence of past experiences on physiology and behaviour, we thought it important to determine if comparing socially naïve and experienced fish would result in different thermal responses. Interestingly, socially experienced fish had a significantly larger T_em_ than socially naïve fish; however, there was no effect on CT_max_. The latter finding may not be surprising since CT_max_ is a physiological marker and, although this variable is plastic (i.e. Morgan *et al*., 2019; Hirakawa & Salinas, 2020), physiology is often less variable than behaviour (Llewelyn *et al*., 2016; Telemeco *et al*., 2009; Huey *et al*., 2003). Our results are also consistent with Currie and Tattersall (2018), where *K. marmoratus* emersion temperatures, but not CT_max_, were influenced by the presence of a mirror. Since socially experienced fish were able to withstand higher temperatures without reducing their TSMs (because CT_max_ was not reduced), they may be using physiological mechanisms that allow them to better cope with temperature elevations. If that is the case, social experience may be particularly important for species that live near their thermal maxima.

The influence of social experience on *pejus* temperature did not appear to be a result of aggressiveness or association time with the social stimulus. However, socially experienced fish had significantly higher levels of whole-body cortisol compared to naïve fish, suggesting that social experience resulted in some physiological stress. Given that fish with higher cortisol concentrations also had larger T_em_, it is a possibility that cortisol is part of the potential mechanism underlying increased T_em_. Notably cortisol has been correlated with TRP expression in a few animal cell models (Ahn *et al*., 2014; Shahsavarani & Perry, 2006; Lin *et al*., 2016). It is tempting to speculate that cortisol may play a role in modulating the sensitivity of TRP thermal receptors in fish (see below).

Surprisingly, the difference in *pejus* temperature between socially experienced and naïve fish disappeared when fish could interact with their mirror reflections or see and smell live conspecifics during thermal stress. Through video analysis, we confirmed that this finding was not due to socially experienced fish displaying increased interest or aggression towards the mirror. As with mirrors, fish did not express higher aggression towards live conspecifics whether they were socially experienced or not. However, socially naïve fish spent more time with conspecifics than socially experienced fish. There are two possible competing hypotheses explaining why the effect of social experience was lost with an acute social interaction: 1) the presence of perceived conspecifics (mirror reflection or live fish) diminishes the stress response of socially experienced fish allowing them to accurately perceive temperature and emerge earlier, or 2) the presence of perceived conspecifics is stressful for the socially naïve fish and the resulting stress response is preventing them from perceiving temperature accurately, thus delaying their emersion. If the second hypothesis is supported, we would expect both socially naïve and experienced fish faced with conspecifics (real or perceived) to have a higher *pejus* temperature than control fish. However, this is not what we observed. Therefore, it seems more likely that the presence of perceived conspecifics is lowering the *pejus* temperature of socially experienced fish, supporting the first hypothesis. Although we know that cortisol levels are higher in socially experienced fish, we do not know if cortisol levels changed with acute social interaction, and this was beyond the scope of our study. It is tempting to speculate that we are observing social buffering where the presence of social partners reduces the stress response of an individual (Hennessy *et al*., 2009; Kiyokawa *et al*., 2007; Hostinar *et al*., 2013).

Social buffering appears to be evolutionarily conserved (Oliveira & Faustino, 2017) as it has been observed in primates (Wittig *et al*., 2016), rodents (Kiyokawa *et al*., 2004), birds (Edgar *et al*., 2015), fish (Culbert *et al*., 2019; Faustino *et al*., 2017; Yusishen *et al*., 2020) and invertebrates (Tian *et al*., 2017). This phenomenon has been identified in a variety of contexts where the duration of the stressor varied from acute to chronic, social partners were present prior, during (coined exposure-type social buffering) or after the stressor (Hennessy *et al*., 2009) and response variables studied were physiological, neural and/or behavioural (Kiyokawa & Hennessy, 2018). Although social buffering is most commonly observed when individuals share a strong bond (e.g., mother – infant, pair-bonded mates) (Kiyokawa & Hennessy, 2018), it can occur between unfamiliar individuals without attachments. For example, zebrafish benefited from new social groups because they had reduced cortisol concentrations (Culbert *et al*., 2019). Although we did not set out to investigate social buffering, it is possible that we are observing a stress-mediating effect of conspecifics here. If perceived conspecifics are reducing emersion temperatures through reduced physiological stress, it may be that social interactions during high temperature events is beneficial to socially experienced fish as otherwise their emersion temperature would dangerously approach their CT_max_. Because the relationship between emersion temperature, sociality and cortisol is not yet established, we do not know whether delayed emersion is related to stress or if it is an appropriate variable to study social buffering. Nevertheless, it remains an interesting and unexpected result.

It should be noted that we observed similar responses in socially experienced fish regardless of whether they were exposed to their mirror reflections or conspecifics. Mirrors are commonly used as a social stimulus with vertebrates that are not capable of self-recognition, including fish, to study behaviours towards conspecifics (Li *et al*., 2018; Earley *et al*., 2000). In some fish species including *K. marmoratus*, an individual’s reflection can evoke greater aggression responses than live conspecifics (Baenninger 1966; Dore *et al*., 1978; Earley *et al*., 2000). However, mangrove rivulus do not interact with mirrors in exactly the same way as with live conspecifics (Li *et al*., 2018). For example, these fish spent less time interacting with mirrors, launch fewer attacks, and switch orientation more frequently than with live conspecifics (Li *et al*., 2018). These muted social response towards mirror reflections have been linked to differences in immediate early gene expression revealing that mirror reflections activate the neural social decision-making network differently than live conspecifics do (Clayton *et al*., 2019; Weitekamp & Hofmann, 2017; Li *et al*., 2018). Therefore, if instead of providing a purely social stimulation as we intended, mirror reflections were perceived as being unfamiliar and frightening (Li *et al*., 2018), it is surprising that that they masked the effect of social experience like live conspecifics. Despite mirrors eliciting different behaviour compared to live conspecifics, we did find that socially naïve fish with mirrors displayed more aggression than socially naïve control fish confirming that mirrors did elicit responses from our laboratory fish.

Our overarching hypothesis was that the delayed emersion of socially experienced fish was the result of desensitized TRPV1 thermoreceptors. In support of this, socially experienced fish emerged at almost twice the capsaicin concentration as socially naive fish suggesting that there is likely a relationship between social experience and thermosensation through the TRPV1 channel. We know that individuals can preferentially choose being social over responding to temperature (i.e. Currie & Tattersall, 2018; Nay *et al*., 2021). Perhaps this trade-off is driven by modified thermosensation of TRPV1. If social experience impacts how fish perceive temperature, the “master abiotic factor” for ectotherms (Brett, 1971), and influences their habitat use (e.g. emerging vs staying submerged), sociality could be considerably more important to survival than previously thought. To our knowledge, there are no studies linking social behaviour to TRP channel activation in any animal. Given that TRP channels are highly conserved among vertebrates (Saito *et al*., 2011), a relationship between sociality and TRP channels could potentially influence behaviour not only in fish, but also in other vertebrates. Notable, insects also share TRP channels with vertebrates (Wei *et al*., 2015) and might then exhibit social behaviour modulated by TRP channels. However, TRP channels can differ biophysically between species (see varying TRPA1 activation temperatures in rodents, reptiles and humans; Story *et al*., 2003; Gracheva *et al*., 2010; Chen *et al*., 2013) and so caution is required when generalizing across species (Garcí-Ávila & Islas, 2019). It is also important to note that TRPV1 is not the only thermoreceptor of fish. TRPV4 is also sensitive to temperature (Boltana *et al*., 2018) and that the number and types of thermoreceptors differs between classes of vertebrates (Nisembaum *et al*., 2015; Boltana *et al*., 2018). Studies specifically looking at TRP channel expression and activation in different species, including fishes, would provide important information to directly establish a link between sociality and thermosensation and help determine if this phenomena is conserved across taxa.

Given that socially experienced fish exhibited higher cortisol levels than their socially naïve counterparts, it is tempting to speculate a mechanistic link between TRPV1 activation and cortisol. Indeed, a few studies have found correlations between cortisol concentration and fish gill TRPV5 mRNA expression (*Oreochromis mossambicus*, Lin *et al*., 2016; *Oncorhynchus mykiss*, Shahsavarani & Perry, 2006) and one study showed that cortisol induced the degradation of TRPV1 receptors in HeLa cells (Ahn *et al*., 2014). Further studies will be necessary to determine if this physiological stress response regulates TRP activation/expression in fish.

In conclusion, we show that social context affected the thermal response of mangrove rivulus to high temperatures. Notably, the effect of social context depended on the source of the social stimulation and the social experience of the individual. We demonstrated that social experience influenced how fish perceived temperature causing them to expose themselves longer to critically warm water. Socially experienced fish emerged at significantly higher capsaicin concentrations than socially naïve fish suggesting that TRPV1 channels of socially experienced fish are less responsive. At this point, we do not know if the decreased thermosensation was through a desensitization of TRPV1 channel itself and/or if socially experienced fish have fewer TRPV1 channels. TRPV1 seems to be involved, but thermal sensation may be more complex given fish have multiple thermosensitive TRP channels. Since emersion behaviour of mangrove rivulus revealed that sociality affects thermal responses and that social experience may impact how fish perceive temperature, past and present social interactions should be carefully considered when studying thermal responses of aquatic ectotherms. This is especially relevant given that in the next decades species, particularly those living in already-warm and polar areas, will need to cope with temperatures beyond their historical experience (IPCC AR6 WGII report, 2022). Understanding how sociality impacts thermal responses in fishes is crucial to accurately predicting how aquatic animals will cope with climate warming.

